# Interpreting short tandem repeat variations in humans using mutational constraint

**DOI:** 10.1101/092734

**Authors:** Melissa Gymrek, Thomas Willems, David Reich, Yaniv Erlich

## Abstract

Identifying regions of the genome that are depleted of mutations can reveal potentially deleterious variants. Short tandem repeats (STRs), also known as microsatellites, are among the largest contributors of *de novo* mutations in humans and are implicated in a variety of human disorders. However, because of the challenges STRs pose to bioinformatics tools, per-locus studies of STR mutations have been limited to highly ascertained panels of several dozen loci. Here, we harnessed bioinformatics tools and a novel analytical framework to estimate mutation parameters for each STR in the human genome by correlating STR genotypes with local sequence heterozygosity. We applied our method to obtain robust estimates of the impact of local sequence features on mutation parameters and used this to create a framework for measuring constraint at STRs by comparing observed vs. expected mutation rates. Constraint scores identified known pathogenic variants with early onset effects. Our constraint metrics will provide a valuable tool for prioritizing pathogenic STRs in medical genetics studies.

## Introduction

Mutations that have negative fitness consequences tend to be eliminated from the population. Thus, identifying regions of the genome that are depleted of mutations has proven a useful strategy for interpreting the significance of *de novo* variation in developmental disorders^1^, prioritizing rare disease variants^2^, and identifying genes or non-coding regions of the genome that are under selective constraint^3,4^. The key idea of these approaches is that mutations occurring at sites evolving under a neutral model are likely to have little effect on reproductive fitness, whereas mutations at intolerant sites are more likely to be involved in severe early-onset disorders.

So far, the genetics community has developed a multitude of methods to assess genetic constraint. These studies have highlighted the importance of a carefully calibrated model of the background mutation process to establish a neutral expectation. For instance, Samocha *et al.^1^* determine the expected number of *de novo* variants per gene based on a neutral model obtained by counting mutations for each possible trinucleotide context in intergenic SNPs. In a different approach, fitCons3 aggregates non-coding regions with similar functional annotations and compares observed variation in those regions to an expectation obtained from presumably neutral flanking regions. Notably, these methods have mainly focused on single nucleotide polymorphisms (SNPs) and to a lesser extent on small indels. As of today, computational methods to analyze and assess the functional impact of repetitive elements in the genome are lacking. Thus, repeat variants are commonly excluded from medical genetics analyses.

To expand the range of interpretation tools to repeat elements, we focused on short tandem repeats (STRs), also known as microsatellites, in the human genome. STRs consist of repeated motifs of 1-6bp and represent about 1.6 million loci^5^, rendering them one of the largest repeat classes. STR mutations are responsible for over 30 Mendelian disorders^6^, many of which are thought to arise spontaneously from *de novo* mutations^7,8^. Emerging evidence suggests STRs play an important role in complex traits^9^ such as gene expression^10^ and DNA methylation11. In addition, analyses of cancer cell lines have shown that STR instability is a chief clinical sign for tumor prognosis^12^, but the functional impact of these instabilities is largely unknown. Evaluating genetic constraint requires two fundamental components: an accurate mutation model and a deep catalog of existing variation. Both of these have been difficult to obtain for repetitive regions of the genome. Current knowledge of the STR mutation process is based on low-throughput studies focusing on an ascertained panel of loci that are highly polymorphic. These include genealogical STRs on the Y chromosome^13,14^, approximately a dozen autosomal STRs from the CODIS (Combined DNA Index System) set used in forensics, and several thousand STRs historically used for linkage analysis^15^. These studies suggest an average mutation rate of approximately 10^-3^ to 10^-4^ mutations per generation^13–17^. However, these loci likely have significantly higher mutation rates than most STRs. Moreover, well characterized STRs consist almost entirely of tetra- or di- nucleotide repeats, which may mutate with different rates and processes compared to other repeat classes. Finally, STR mutation rate studies have been based on small numbers of families and show substantial differences regarding absolute mutation rates and their patterns (**Supplemental Table 1**).

Here, we developed a framework to measure constraint at individual STRs that benefits from a novel method to obtain observed and expected mutation rates at each locus. We developed a robust quantitative model that harnesses population-scale genomic data to estimate locus-specific mutation dynamics at each STR by correlating local SNP heterozygosity with STR variation. After extensive validation, we applied this model to estimate mutation rates at more than one million STRs using whole genome sequencing of 300 unrelated samples from diverse populations^18^. Using these results, we built a model to predict mutation parameters from local sequence features and measured constraint at each STR locus. One caveat is that our method is primarily applicable to STRs that can be completely spanned by short reads, and does not accurately describe large expansion mutations observed in conditions such as Huntington’s Disease or Fragile X Syndrome. We show that our constraint metric can be used to predict clinical relevance of individual STRs, including those in genes with known implications in developmental disorders. This framework will likely enable better assessment of the role of STRs in human traits and will inform future work incorporating STRs into human genetics studies.

## Results

### A method to estimate local mutation parameters

We first sought to develop a method to estimate mutation parameters at each STR in the genome by fitting a model of STR evolution to population-scale data. A primary requirement of our method is a model of the STR mutation process that fits observed variation patterns. Motivated by the poor fit of the widely used generalized stepwise mutation model (GSM) to our data (**Supplemental Note 1**), we developed a novel length-biased version of GSM that closely recapitulates observed population-wide trends (**Supplemental Notes 2,3; Supplemental Figures 1,2**), including a saturation of the STR molecular clock over time. Our model includes three parameters: *μ* denotes the per-generation mutation rate, *β* describes the strength of the directional bias of mutation, and *p* describes the geometric mutation step size distribution. Recently, we developed a method called MUTEA that employs a similar model to precisely estimate individual mutation rates for Y chromosome STRs (Y-STRs) from population-scale sequencing panels of unrelated individuals. MUTEA models STR evolution on the underlying SNP-based Y phylogeny^19^. We found good concordance (r^2^=0.87) between MUTEA and traditional trio-based methods and high reproducibility (r^2^=0.92) across independent datasets. However, the main limitation of this approach is that it requires full knowledge of the underlying haplotype genealogy, which is difficult to obtain for autosomal loci.

To analyze the mutation rates of autosomal STRs, we extended MUTEA to analyze pairs of haplotypes. The key insight of our mutation rate estimation procedure is that different classes of mutations provide orthogonal molecular clocks (**Figure 1**). Consider a pair of haplotypes consisting of an STR and surrounding sequence. The SNP heterozygosity is a function of the time to the most recent common ancestor (TMRCA) of the haplotypes and the SNP mutation rate. On the other hand, the squared difference between the numbers of repeats of the two STR alleles (defined as the allele squared distance, or ASD) is a separate function of the TMRCA. The distribution of ASD values observed for a given TMRCA is determined by our mutation model. Using known parameters of the SNP mutation process, we can measure the local TMRCA to calibrate the STR molecular clock^15^.

**Figure 1:**
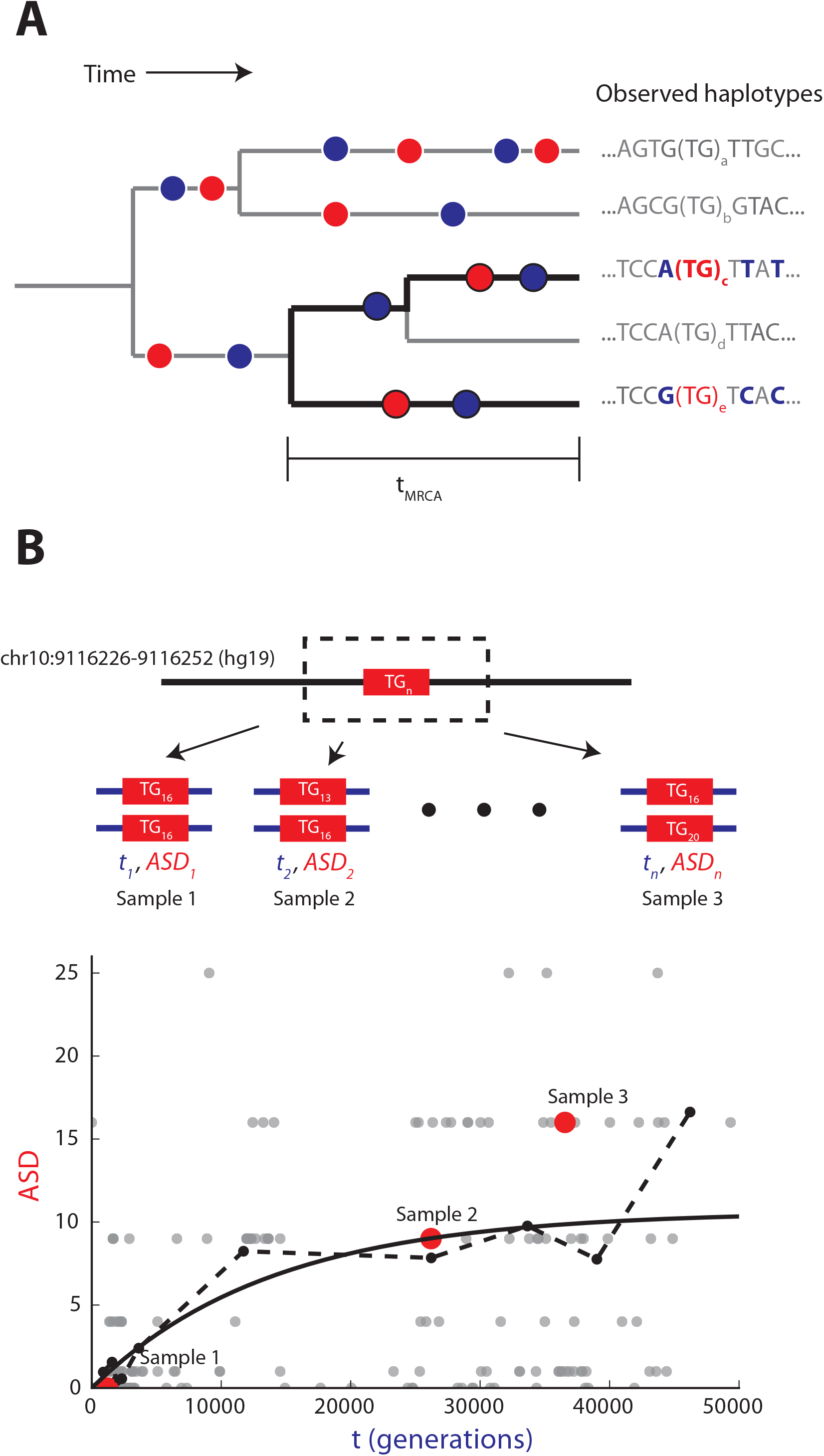
Estimating STR mutation parameters from diploid data. (A) SNPs and STRs give orthogonal molecular clocks. The tree represents an example evolutionary history of an STR locus. Red dots denote STR mutation events. Blue dots represent SNP mutation events. Black branches denote an observed diploid locus, consisting of two haplotypes from the tree. **(B) Correlating local TMRCA with STR genotypes allows per-locus mutation rate estimation.** For each diploid STR call, we use SNP heterozygosity to extract the TMRCA (blue) of the surrounding region and the squared length difference between STR alleles (ASD, in red). Our STR mutation model describes the expected ASD for a given TMRCA (solid black line). Gray dots give data points for each sample, red dots represent three example samples, and the dashed black line gives the sliding window mean.

Our method takes as input unphased STR and SNP genotypes and returns maximum likelihood estimates of STR mutation parameters. The TMRCA is approximated by local SNP heterozygosity using a pairwise sequentially Markovian coalescent model^20^ (**Methods**). ASD is calculated directly from a diploid STR genotype as the squared difference in the number of repeats of each allele. Our maximum likelihood framework allows us to estimate parameters at a single STR or jointly across many loci. A potential caveat is that haplotype pairs may have shared evolutionary history and thus are not statistically independent, which is not expected to bias our estimates but will artificially shrink standard errors. To account for this non-independence, we adjust standard errors by calibrating to ground truth simulated and capillary electrophoresis datasets (**Supplemental Note 4, Supplemental Figure 3**).

### Validating parameter estimates

We first evaluated our estimation procedure on STR and SNP genotypes simulated on haplotype trees using a wide range of mutation parameters. To evaluate our method on unphased diploid data, we formed a set of 300 “diploids” by randomly selecting leaf pairs and recording the TMRCA and STR allele lengths. To test the effects of genotyping errors, we simulated “stutter” errors using the model described in Willems *et al*.^19^ and used the expectation-maximization framework we developed previously^21^ to estimate per-locus stutter noise and correct for STR genotyping errors.

Our method obtained accurate per-locus estimates for *μ* for most biologically relevant parameter ranges (**Figure 2A**). Notably, estimates for *p* and *β* were less precise (**Supplemental Figure 4**) and thus downstream analyses focused on mutation rates. The main limitation of our method is an inability to capture low mutation rates. Informative estimates could be obtained for rates >10^-6^. This presumably stems from the low number of total mutations observed (median 1 mutation for μ=10^-6^ in 300 samples). Aggregating loci, or equivalently analyzing larger sample sizes, gives higher power to estimate low mutation rates due to the higher number of total mutations observed. By analyzing loci jointly, we could accurately estimate mutation rates down to 10^-6^ with 30 or more loci and 10^-7^ with 70 or more loci (**Figure 2B**). As expected, inferring and modeling stutter errors correctly removed biases induced by stutter errors (**Supplemental Figure 5**).

**Figure 2:**
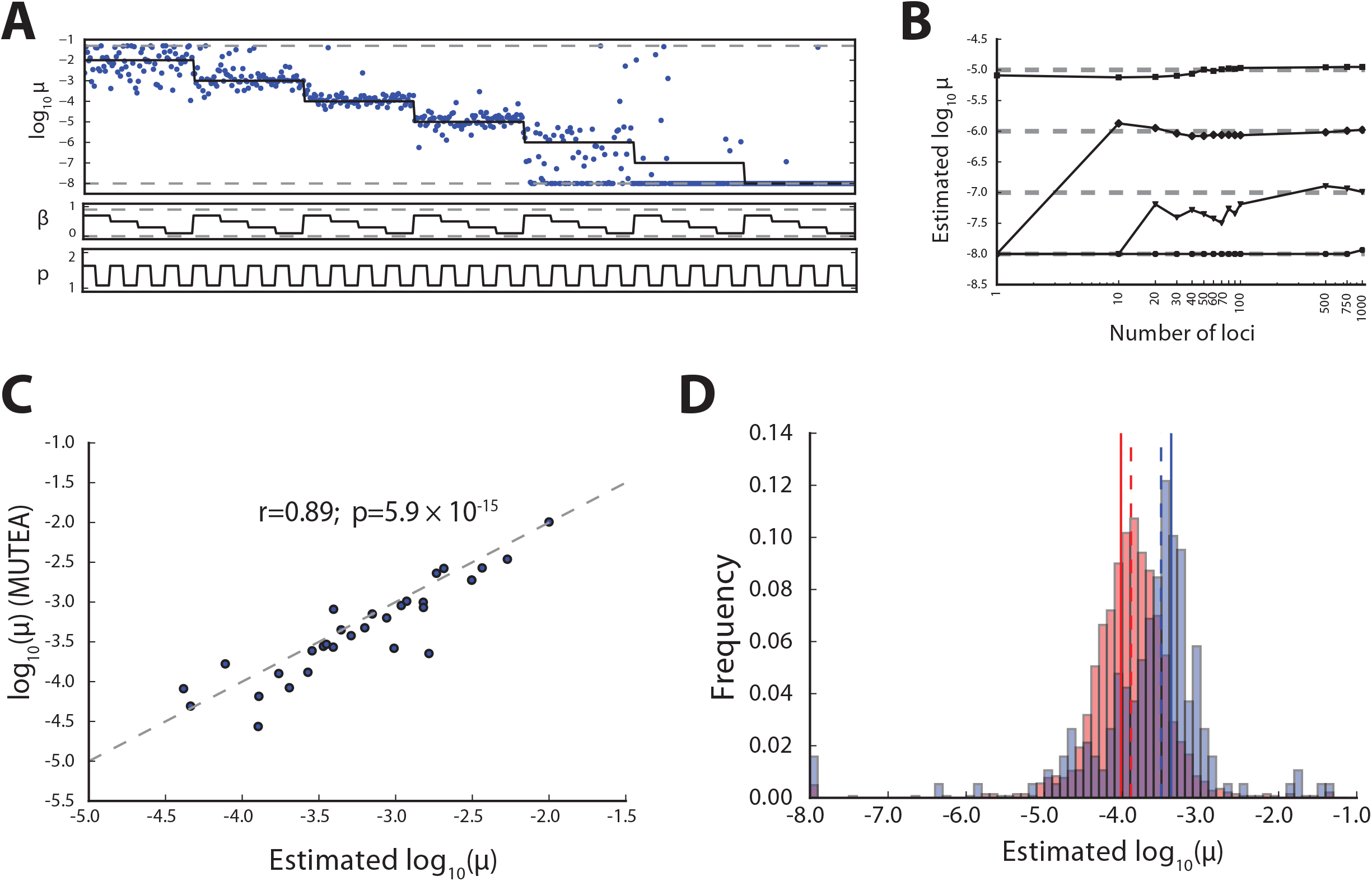
Accurate estimation of STR mutation parameters from simulated data. (A) Per-locus estimates of mutation rate. Solid black lines give simulated values. Blue dots give per-locus estimates. Dashed gray lines give boundaries enforced during numerical optimization. **(B) Jointly estimating parameters across loci allows inference of slow mutation rates.** Black lines give joint estimates for different simulated mutation rates (circles=10^-8^, triangles=10^-7^, diamonds=10^-6^, squares=10^-5^). Dashed gray lines give simulated values. **(C) Y-STR mutation rate parameters are concordant across estimation methods.** Mutation rate estimates from this study compared to those returned by MUTEA. Gray dashed lines denote the diagonal. **(D) Autosomal mutation rate estimates are concordant with *de novo* studies.** Dashed lines give median estimate across loci. Solid lines give empirical mutation rate from trio data analyzed by Sun *et al.* Red=dinucleotides; blue=tetranucleotides.

We next evaluated the ability of our method to obtain mutation rates from population-scale sequencing of Y-STRs whose mutation rates have been previously characterized. We analyzed 143 males sequenced to 30-50x by the Simons Genome Diversity Project^18^ (SGDP) and 1,243 males sequenced to 4-6x by the 1000 Genomes Project^22^. We used all pairs of haploid Y chromosomes as input to our maximum likelihood framework. We compared our results to two orthogonal mutation rate estimates: our previous MUTEA method19 and a study that examined 500 father-son duos^13^. We found that the mutation rate estimates were consistent across sequencing datasets (r=0.90; p=1.5×10^-18^; n=48) (**Supplemental Figure 6**). Encouragingly, our rate estimates were similar to those reported by MUTEA on the SGDP dataset (r=0.89; p=5.9×10^-15^; n=41) (**Figure 2C**). Furthermore, our estimates were significantly correlated with those reported by Ballantyne *et al.* (r=0.78; p=2.0×10^-9^; n=41) (**Supplemental Figure 6**), a substantial improvement over results obtained using a traditional stepwise mutation model (r=0.37; p=0.0150; n=41), validating our choice of mutation model.

Finally, we evaluated our method on a subset of well characterized autosomal diploid loci. We first analyzed the forensics CODIS markers, which have well-characterized mutation rates estimated across more than a million meiosis events (http://www.cstl.nist.gov/strbase/mutation.htm). Mutation rates were concordant with published CODIS rates (r=0.90, p=0.00016, n=11) (**Supplemental Figure 7**). We also compared to di- and tetranucleotide mutation rates previously estimated by Sun *et al.* by aggregating data from 1,634 loci in 85,289 Icelanders^15^. Mutation rates were in strong agreement (**Figure 2D; Supplemental Figure 8**), which is especially encouraging given that the Sun *et al.* STR genotypes were obtained using an orthogonal method of capillary electrophoresis.

### Characterizing the STR mutation process using diverse whole genomes

Next, we applied our mutation rate estimation method genome-wide. We analyzed 300 individuals from diverse genetic backgrounds sequenced to 30-50x coverage by the SGDP Project^18^. We aligned reads to the hg19 reference genome using BWA-MEM^23^ and the resulting alignments were used as input into lobSTR^24^ (**Methods**). High quality SNP genotypes were obtained from our previous study^18^. We used these as input to PSMC^20^ to estimate the local TMRCA between haplotypes of each diploid individual. For each locus, we adjusted genotypes for stutter errors (**Supplemental Figure 9; Supplemental Table 2**, **Methods**) and used adjusted genotypes as input to our mutation rate estimation technique. After filtering (**Methods**), 1,251,510 STR loci with an average of 249 calls/locus remained for analysis (**Supplemental Dataset 1**). Results were concordant with mutation rates predicted by extrapolating MUTEA to autosomal loci (r=0.71; p<10-16; n=480,623) (**Supplemental Figure 10**), suggesting that our mutation rate estimation is robust even in the case of unphased genotype data from modest sample sizes.

Per-locus mutation rates for each repeat motif length varied over several orders of magnitude, ranging from 10^-8^ to 10^-2^ mutations per locus per generation (**Supplemental Figure 11; Supplemental Table 3**). Median mutation rates were highest for homopolymer loci (log_10_μ=-5.0) and decreased with the length of the repeat motif, with most pentanucleotides and hexanucleotides below our detection threshold. Interestingly, homopolymers also showed markedly higher length constraint compared to other loci, suggesting an increased pressure to maintain specific lengths. Step size distributions also differed by repeat motif length. Homopolymers (median p=1.00) and to a lesser extent repeats with motif lengths 3-6 (median p=0.95) almost always mutate by a single repeat unit. On the other hand, dinucleotides are more likely to mutate by multiple units at once, consistent with previous studies^15^. Overall, our results highlight the diverse set of influences on the STR mutation process, and suggest there is limited utility to citing a single set of STR mutation parameters.

### A framework for measuring STR constraint

Encouraged by the accuracy of our per-locus autosomal parameter estimates, we sought to create a framework to evaluate genetic constraint at STRs by comparing observed to expected mutation rates. Our framework relies on generating robust predictions of per-locus mutation rates based on local sequence features and comparing the departure of the observed rates from this expectation (**Figure 3A**). STRs whose observed mutation rates are far lower than expected are assumed to be under selective constraint, and thus more likely to have negative fitness consequences.

**Figure 3:**
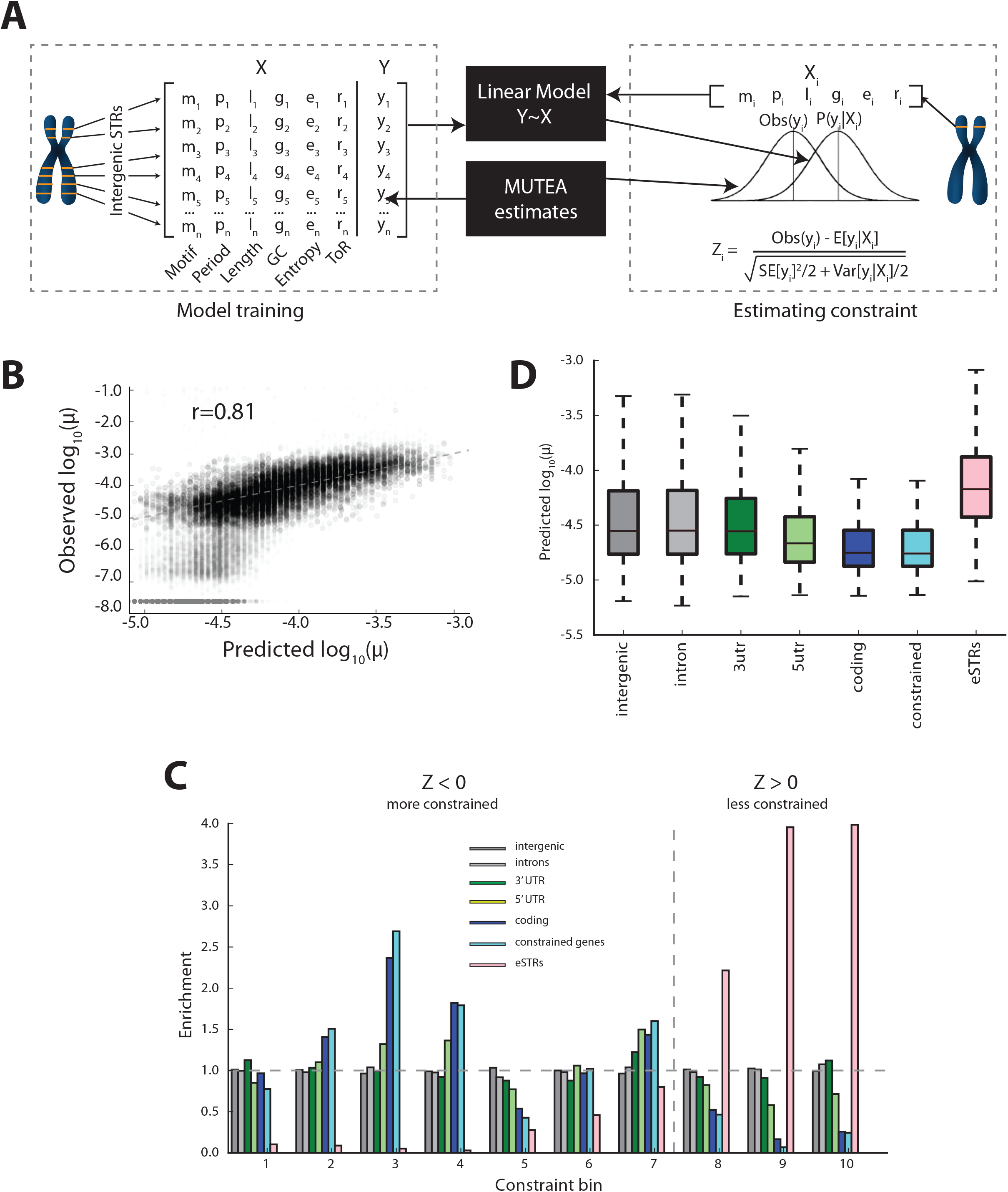
A framework for measuring STR constraint. (A) Schematic of constraint framework. In the model training phase, a linear model is trained to predict mutation rates from local sequence features. In the estimation phase, constraint is measured by comparing predicted mutation rates to observed rates. **(B) Sequence features are predictive of mutation rate**. Comparison of predicted vs. observed mutation rates for a held out test set of intergenic loci. Gray dots denote loci with high or undefined standard errors that were excluded from model training. **(C) Enrichment of gene annotations by constraint bin.** X-axis gives bins defined by Z-score deciles. Y-axis gives the fold enrichment of each annotation in each bin. The dashed line gives the boundary between constrained (Z<0) and non-constrained (Z>=0) scores. **(D) Predicted mutation rates by annotation.** For **(C)** and **(D),** constrained denotes genes with missense constraint score >3 as reported by ExAC.

We began by evaluating whether local sequence features can accurately predict STR mutation rates. We examined the relationship between STR mutation rate and a variety of features, including total STR length, motif length, replication timing, and motif sequence (**Supplemental Figure 12**). While all features were correlated with mutation rate (**Supplemental Table 4**), total uninterrupted repeat sequence length and motif length were by far the strongest predictors, as has been previously reported by many studies^19,25^. These features were combined into a linear regression model to predict per-locus mutation rates. We stringently filtered the training data to consist of presumably neutral (intergenic) loci with the best model performance. Analysis was restricted to STRs with motif lengths of 2-4bp with reference length ≥20bp and small standard errors (**Methods**), since this subset showed mutation rates primarily in the range that our model can detect. Using this filtered set of markers, a linear model explained 65% of variation in mutation rates in an independent validation set (**Figure 3B**).

We next developed a metric to quantify constraint at each STR by comparing observed to expected mutation rates (**Supplemental Dataset 1**). Our constraint metric is calculated as a Z-score, taking into account errors in both the predicted and observed values (**Methods**). Negative Z-scores denote loci that are more constrained than expected, and vice versa. Constraint scores for loci with detectable mutation rates followed the expected standard normal distribution (**Supplemental Figure 13**). However, loci with mutation rates below our detection threshold of 10^-6^ do not have reliable standard error estimates and had downward biased scores. Nevertheless, these loci are informative of a constraint signal for instances where the predicted mutation rate is high but the observed rate is below our detection threshold. Thus, rather than analyzing distributions of raw constraint scores, we binned scores by deciles and examined enrichments for functional annotations in each bin. For comparison, we also calculated mutation rates and constraint scores assuming a generalized stepwise model (**Methods**) and found that mutation rates and constraint scores were similar (r=0.88 and r=0.56 for mutation rates and constraint scores, respectively). All constraint scores analyzed below were calculated using the length-constrained model.

### STR constraint scores give insights into human phenotypes

Observed Z-scores are concordant with biological expectations across genomic features. Introns, intergenic, and 3’-UTR regions closely matched neutral expectation (**Figure 3C**). On the other hand, STRs in coding exons showed significantly reduced mutation rates compared to the null model. These trends were recapitulated in the expected mutation rates (**Figure 3D**), suggesting that STRs under constraint are also under evolutionary pressure to maintain sequence features contributing to lower mutability. Additional analysis of STR constraint in coding regions is given in **Supplemental Note 5** and **Supplemental Figure 14**. In contrast to strong levels of constraint in coding exons, the STRs that we had previously identified to act as expression quantitative trait loci (eQTLs)^10^ showed a marked lack of constraint, consistent with previous observations in the Exome Aggregation Consortium (ExAC) dataset^26^ showing highly constrained genes are depleted for eQTLs.

Constraint can provide a useful metric to prioritize potential pathogenic variants and interpret the role of individual loci in human conditions. Notably, this metric is most sensitive to early-onset disorders, as mutations involved in later onset disorders generally do not affect fitness and are thus expected to follow neutral patterns. Additionally, constraint is most sensitive to deleterious mutations following dominant inheritance patterns, since recessive mutations are eliminated at much slower rates. Consistent with this theory, STRs implicated in early onset dominant diseases show significantly higher constraint than expected (**Figure 4**). We focused on STRs that can be genotyped from high throughput sequencing data and are involved in congenital disorders. Notably, this excludes most large repeat expansions such as those involved in Huntington’s Disease or Fragile X Syndrome. First, we examined polyalanine and polyglutamine tracts in *RUNX2.* Even mild expansion of four glutamine residues has been shown to result in congenital cleidocranial dysplasia (OMIM: 119600)^27,28^. Both repeats showed constrained mutation rates, with the polyglutamine repeat in the most constrained bin (Z=-11.3). Next, we tested a polyalanine expansion in *HOXD13*, which causes a severe form of synpolydactyly (OMIM: 186000). Again, a mild expansion (7 additional residues) has been shown to be pathogenic^29^. This repeat was on the boundary of the most severe constraint bin (Z=-10.9). As a negative control, we also tested constraint at the CODIS loci used in forensics, which have been specifically ascertained for their high polymorphism rates and are likely neutral. As expected, the CODIS markers have weak constraint scores, and exhibit slightly higher mutation rates than expected (Z>0) (**Figure 4**).

**Figure 4:**
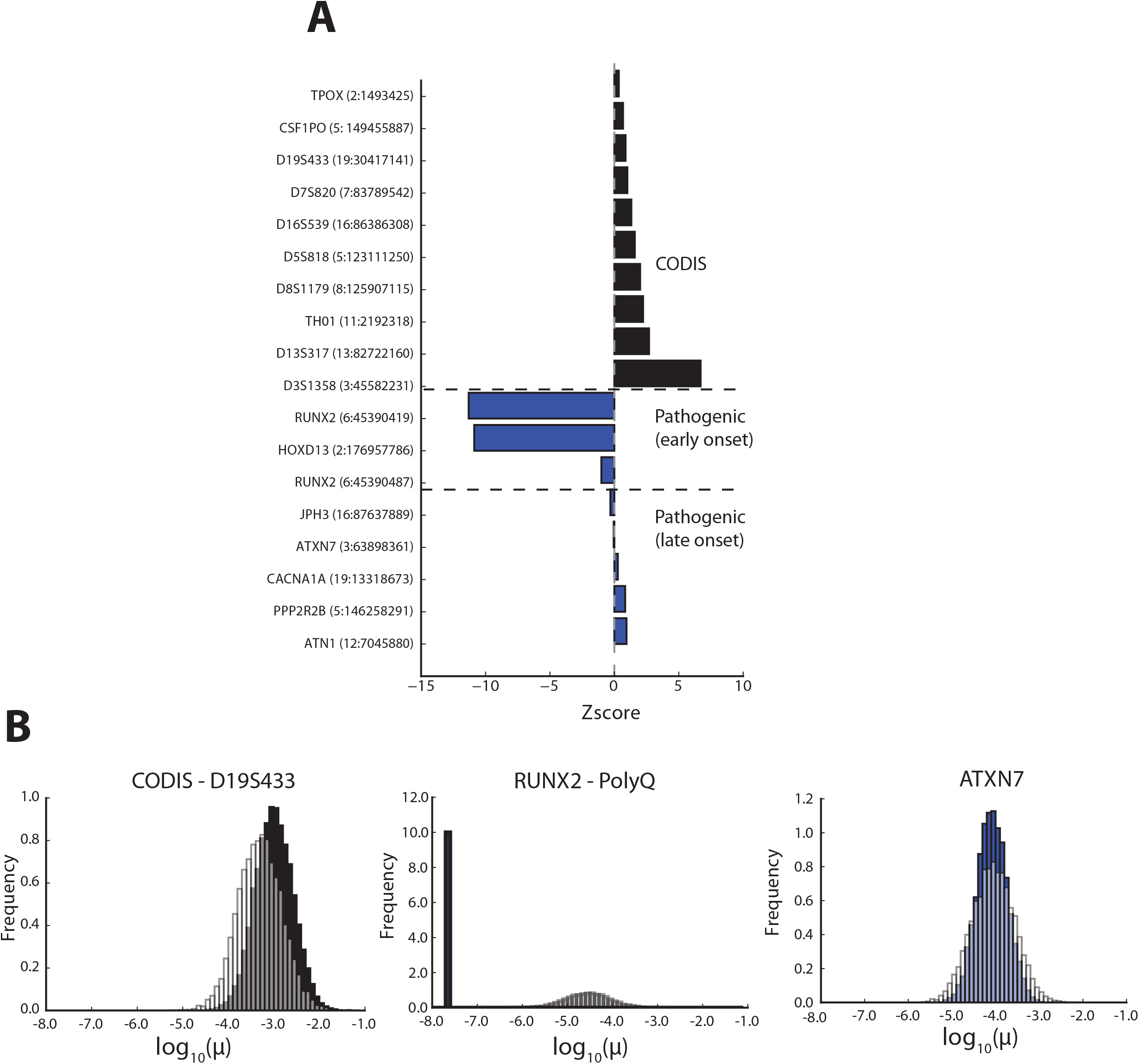
Constraint scores can be used for STR prioritization. (A) Z-scores for example loci. Black gives CODIS forensics markers. Blue give known pathogenic STRs. For each STR, the CODIS marker or gene name is given, and the chromosomal location (hg19) is indicated in parentheses. **(B) Example distributions of observed vs. expected mutation rates.** The left panel shows a CODIS STR (D19S433), a presumably neutral STR. The middle panel shows a highly constrained polyglutamine repeat in *RUNX2* for which a mild expansion is implicated in an early onset disorder. The right panel shows a polyglutamine repeat in *ATXN7*, implicated in a late onset disorder and accordingly not highly constrained. White bars=expected mutation rate distribution. Solid bars=observed mutation rate distribution.

More broadly, we found STRs are highly enriched in genes that are involved in developmental processes (p=9.78×10^-38^). Consistent with this result, three of the ten most highly constrained coding STRs in our dataset are in genes with previously reported developmental disorders following autosomal dominant inheritance patterns that have yet to be associated with pathogenic STRs: *GATA6* (congenital heart defects, OMIM: 600001), *SOX11* (mental retardation, OMIM: 615866), and *BCL11B* (Immunodeficiency 49, OMIM: 617237) (**Supplemental Table 5**). On the other hand, we found that pathogenic STRs of late onset STR expansions disorders such as cerebellar ataxias were not highly constrained, and showed mutation rates very close to predicted values (**Figure 4**). These disorders often do not occur until the fourth or fifth decade of life^30^, and thus are not expected to be under strong purifying selection. Taken together, these results suggest STR constraint scores will provide a useful metric by which to prioritize rare pathogenic variants involved in severe developmental disorders.

To facilitate use by the genomics community, genome-wide results of our mutational constraint analysis are provided in **Supplemental Dataset 1**, which can be analyzed with standard genomics tools such as BEDtools^31^.

## Discussion

Metrics for quantifying genetic constraint by comparing observed to expected variation have provided a valuable lens to interpret the impact of *de novo* SNP variants. These have been widely used for applications including quantifying the burden of *de novo* variation in neurodevelopmental disorders^1,32^, identifying individual genes constrained for missense or loss of function variation^26^, and more recently to measure constraint in non-coding elements^4,33^. However, the mutation rate at SNPs is sufficiently low that any given nucleotide has a low probability of being covered by a polymorphism even in very large datasets of human variation (e.g. a dataset of more than 60,000 exomes contained about 1 polymorphism per 8 nucleotides^26^). Thus, the information provided by SNP variation is never sufficient to provide a direct measurement of the likely evolutionary constraint on a particular mutation. In contrast, the much higher mutation rate in STRs makes it possible to precisely measure constraint on a per-site basis even with as few as 300 whole genomes.

We combined a deep catalog of STR variation^18^ with a novel model of the STR mutation process to develop an accurate method for measuring per-locus STR mutation parameters by correlating STR variation with local sequence heterozygosity across haplotype pairs. We used this method to estimate mutation rates at more than 1 million individual STRs in the genome. Observed STR mutation rates vary over several orders of magnitude, suggesting it is not useful to cite a single mutation rate for all STRs. Median genome-wide mutation rates were far lower than previously reported^16,17,25,34^. This is consistent with the fact that most well studied STR panels were specifically ascertained for their high heterozygosity, needed for traditional STR applications such as forensics or genetic linkage analysis. Our estimates confirm many known trends in STR mutation, such as the dependence of mutation rate on total STR length and the tendency of dinucleotide repeats to mutate in larger units than tetranucleotides^25^. Moreover, this large dataset allows us to exclude the possibility that certain sequence features such as local GC content play a strong role in determining STR mutation rates.

We showed that by comparing observed to expected mutation rates, we can measure genetic constraint at individual loci and use our constraint metric to prioritize potentially pathogenic variants. Importantly, our approach provides a biologically agnostic approach to assessing the importance of individual loci, as it relies entirely on observed genetic variation. While our analyses focused on STRs, the framework developed here can be easily extended to any class of repetitive variation for which accurate genotype panels are available. In future studies, we envision this work will provide a much needed framework to interpret the dozens of *de novo* variants at STRs and other repeats arising in each individual, especially in the context of severe early onset disorders. Beyond analyzing *de novo* variation, accurate models of STR mutation will allow scanning for STRs under selection^35^, identifying rapidly mutating markers for forensics or genetic genealogy^19,36^, and enabling improved statistical methods for incorporating STRs into quantitative genetics studies.

Our mutation rate estimation method and constraint metric face several limitations. First, estimating mutation rates in several hundred samples is only accurate for mutation rates down to approximately 10-6. Loci with slower mutation rates produce biased results, limiting our ability to predict and measure mutation rates at a large number of loci, including the majority of protein coding STRs. While we can detect general signals of constraint for slowly mutating STRs, larger sample sizes will allow for more accurate constraint scores and thus more informative prioritization. Second, our method analyzes pairs of haplotypes rather than the entire evolutionary history of a locus. While this has the advantage of allowing estimation across unphased data, it discards valuable information present in the full haplotype tree, and thus limits the scope of models that can be considered. For example, it precludes modeling allele length-specific mutation rates, which requires estimating ancestral states on the full haplotype tree. Finally, there are additional aspects of the STR mutation process not modeled here. Our method focus on short stepwise mutations occurring at relatively stable STRs. Unstable expansions, such as those occurring in trinucleotide repeat disorders, likely mutate by different models. Our model also ignores the effect of sequence interruptions and interaction between alleles, both of which have been hypothesized to influence STR mutation patterns^19,35,37^.

Future bioinformatic advances will likely overcome many of these issues and improve the precision of our estimates. In particular, while our method works on unphased data, phased STR and SNP haplotypes would allow analysis of the entire haplotype tree at a given locus as is done by MUTEA, improving our accuracy and allowing us to consider a broader range of mutation models. Additionally, our current tools are limited to STRs that can be spanned by short reads, and thus exclude many well known pathogenic loci such as those involved in trinucleotide repeat expansion disorders. We envision that long read and synthetic long read technologies will both enable analysis of a broader class of repeats and provide an additional layer of phase information. Finally, larger sample sizes will allow more accurate analysis of constraint for slow-mutating loci. Taken together, these advances will provide a valuable framework for interpreting mutation and selection at hundreds of thousands of STRs in the genome and will help prioritize STR mutations in clinical studies.

## Methods

### STR mutation model

We model STR mutation using a discrete version of the Ornstein-Uhlenbeck process described in detail in **Supplemental Note 2**. Our model assumes STR mutations occur at a rate of μ mutations per locus per generation according to a step-size distribution with first and second moments:

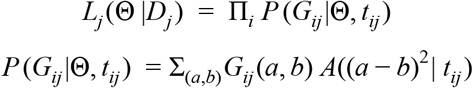

where *a*_*i*_ is the length of the STR allele after mutation *i* and *a*_*i*+1_ is the length after mutation *i* + 1. This implies that long alleles (>0) tend to decrease back toward 0 and short alleles (<0) tend to increase toward 0. For all analyses, all alleles are assumed to be relative to the major allele, which is set to 0.

### Mutation parameter estimation

We extended the MUTEA framework to estimate parameters at diploid loci for which the underlying haplotype tree is unknown. For each sample genotyped at locus j, we obtain t_ij_, the TMRCA between the two haplotypes of sample i, and a distribution G_ij_, where G_ij_(a,b) gives the posterior probability that sample i has genotype (a,b). We initially assume that haplotype pairs are independent and maximize the following likelihood function at locus j:

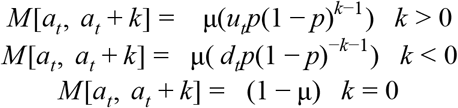

Where Θ = {μ, β, *p*}, *Dj* = {(*G*_1*j*_, *t*_1*j*_), (*G*_2*j*_, *t*_2*j*_)…(*G*_*nj*_, *t*_*nj*_)}, and *A*(*x*|*t*) gives the probability of observing a squared distance of x between alleles on haplotypes with a TMRCA of t. We used the Nelder-Mead algorithm to minimize the negative of the log-likelihood and imposed boundaries of μ ∈ [10^-8^, 0.05], β ∈ [0, 0.9], *p* ∈ [0.7, 1.0].

To compute the function *A*, we first build a transition matrix *M* of size *L* × *L*, where *L* is the number of allowed alleles. *M* [*a*, *b*] gives the probability that allele *a* mutates to allele *b* in a single generation. Step sizes were set based on the model described in **Supplemental Note 2**:

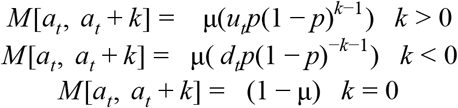

where 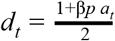 and 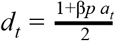.

M represents a stochastic process, and thus *M ^T^* gives transition probabilities along a branch T generations long. A single row *M ^T^* [*a*, :] gives the expected allele frequency spectrum of a locus for which the ancestral allele was *a* and the MRCA was T generations ago. We can use this to derive the probability of observing a given squared distance between two alleles separated by T generations:

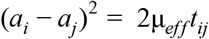

In our data, we do not know the ancestral allele *a* for each pair of haplotypes. However, under our model of STR evolution, *A* does not depend on the ancestral allele and so we assume 0 as the ancestral allele for simplicity. Notably, we have assumed haplotype pairs are statistically independent. While this does not bias our results, standard errors must be adjusted as described in **Supplemental Note 4**.

### Estimating mutation parameters using a generalized stepwise model

Under a generalized stepwise model (GSM), the ASD should be linearly related to the TMRCA between a pair of haplotypes^38^:

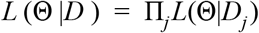

Where a_i_ and a_j_ are the length of STR alleles on two haplotypes i and j, t_ij_ is the TMRCA between that pair of haplotypes, and μ_*eff*_ is the effective mutation rate. Effective mutation rate is defined as μ_*eff*_ = μσ^2^*m*, where μ is the per-generation mutation rate of the locus and step sizes are drawn from a distribution with mean 0 and variance σ^2^*m*.

For each locus, we calculated μ_*eff*_ by regressing ASD on TMRCA and dividing the resulting

### Joint estimation of mutation parameters across multiple loci

The MUTEA approach can be easily extended to estimate mutation parameters in aggregate by jointly maximizing the likelihood across multiple loci at once:

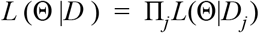

To minimize computation and because β and p tended to be less consistent across loci, we first perform per-locus analyses to obtain individual estimates for β and p. We then hold these parameters constant at the mean value across all loci and only maximize the joint likelihood across μ.

### Simulating SNP-STR haplotypes

We used fastsimcoal^39^ to simulate coalescent trees for 600 haplotypes using an effective population size of 100,000. We then forward-simulated a single STR starting with a root allele of 0 using specified values of μ, β, and. 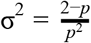. Mutations were generated according to a Poisson process with rate λ = 1/μ and following the model described above. We chose 300 random pairs of haplotypes to form “diploid” individuals to use as input to our estimation method. We simulated reads for each locus assuming 5x sequencing coverage, with each read equally likely to originate from each allele. Stutter errors were simulated using the model described in Willems *et al*._19_ with *u* = 0.1, *d* = 0.05, and ρ_*s*_ = 0.9. This indicates that stutter noise causes the true allele to expand or contract with 10% or 5% frequency, respectively, and that error sizes are geometrically distributed with 10% chance of mutating by more than one repeat unit. For estimating per-locus parameters, we performed 10 simulations with each set of parameters.

### Datasets

#### Previously published mutation rate estimates

MUTEA mutation rate and length bias estimates for the 1000 Genomes dataset were obtained from Table S1 in Willems *et al*.^19^ *De novo* Y-STR mutation rate estimates were obtained from Table S1 of Ballantyne *et al*.^13^ CODIS mutation rates were obtained from http://www.cstl.nist.gov/strbase/mutation.htm.

#### Annotations

Local GC content and sequence entropy were obtained from the “strinfo” file included in the lobSTR hg19 reference bundle. Missense constraint scores were downloaded from the ExAC website http://exac.broadinstitute.org/downloads.

### STR genotyping

#### Profiling STRs from short reads

Raw sequencing reads for the SGDP dataset were aligned using BWA-MEM. Alignments were used as input to the allelotype tool packaged with lobSTR24 version 4.0.2 with non-default flags “—filter-mapq0 –filter-clipped –max-repeats-in-ends 3 –min-read-end-match 10 –dont-include-pl –min-het-freq 0.2 –noweb”. STR genotypes are available on dbVar under accession nstd128. Y-STRs for the 1000 Genomes data were previously profiled^24^ and were preprocessed as described in^19^.

#### Filtering to obtain high quality STR calls

Y-STR calls for SGDP were filtered using the lobSTR_filter_vcf.py script available in the lobSTR download with arguments “--loc-max-ref-length 80 --loc-call-rate 0.8 --loc-log-score 0.8 --loc-cov 3 --call-cov 3 --call-dist-end 20 --call-log-score 0.8” and ignoring female samples. Autosomal samples were filtered using “--loc-max-ref-length 80 --loc-call-rate 0.8 --loc-log-score 0.8 --loc-cov 5 --call-cov 5 --call-dist-end 20 --call-log-score 0.8”.

### Calculating local TMRCA

As described in^18^, we used the pairwise sequential Markovian coalescent (PSMC)^20^ to infer local TMRCAs across the genome in each sample. For each region overlapping an STR, we calculated the geometric mean of the upper and lower heterozygosity estimates returned by PSMC. We scaled heterozygosity to TMRCA based on the genome-wide average PSMC estimate (0.00057) of a French sample with a previously estimated genome-wide average TMRCA of 21,000 generations^25^. To accommodate errors in this scaling process, final mutation rate estimates were scaled to match the mean values of published *de novo* rates (see below).

### Pairwise Y chromosome analysis

TMRCAs for each pair of SGDP Y-chromosomes was calculated using pairwise sequence heterozygosity. We scaled this to TMRCA using the relationship *h*_*i*_/(2μ_*Y SNP*_), where *hi* is the heterozygosity of pair *i* and μ_*Y SNP*_ is the Y-chromosome SNP mutation rate. μ_*Y SNP*_ was set to 2.1775×10^-8^ as reported by Helgason *et al*.^40^ For the 1000 Genomes set, we obtained a Y-phylogeny that was built by the 1000Y analysis group^41^. We scaled the tree using the method described previously^19^. For each dataset, we used pairwise TMRCA and allele squared distance estimates as input to our maximum likelihood procedure.

### Scaling mutation parameters

Our TMRCA estimates, and thus mutation rate estimates, scale linearly with the choice of SNP mutation rate. To account for this and to compare estimates between datasets, we scaled our mutation rates by a constant factor such that the mean STR mutation rates between datasets were identical. Genome-wide estimates are scaled based on the comparison with CODIS rates.

### Measuring STR constraint

#### Predicting mutation rates from local sequence features

We trained a linear model to predict log_10_ mutation rates from local sequence features including GC content, replication timing, sequence entropy, motif sequence, motif length, total STR length, and uninterrupted STR length. The model was built using presumably neutral intergenic loci, with 75% of the loci reserved for training and 25% for testing. While all features were correlated with mutation rates, the best test performance was achieved using only motif length and uninterrupted STR length. Models were built using the python statsmodels package (http://www.statsmodels.org/).

Model training was restricted to STRs whose mutation rates could be reliably estimated. We filtered STRs with total length <20bp, since the majority of shorter STRs returned biased mutation rates at the optimization boundary of 10^-8^. We further filtered STRs with standard errors equal to 0, >0.1, or undefined (usually indicating the lower optimization boundary of 10^-8^ was reached). However, these loci were included in testing and in downstream analysis as the majority of coding STRs fell into this category.

#### Calculating Z-scores

Constraint scores are calculated for each locus *i* as:

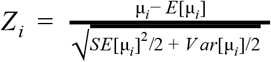

Where μ_*i*_ is the observed mutation rate, *SE*[μ_*i*_] is the standard error of the observed mutation rate, *E*[μ_*i*_] is the predicted mutation rate, and *V ar*[μ_*i*_] is the variance of the prediction. In all cases, μ_*i*_ refers to the log10 mutation rate, with the log_10_ notation omitted for simplicity.

#### Constraint score analysis

GO analysis was performed using goatools (https://github.com/tanghaibao/goatools). OMIM disease annotations were accessed on December 8, 2016.

### Data availability

**Supplemental Dataset 1** is available at https://s3-us-west-2.amazonaws.com/strconstraint/Gymrek_etal_SupplementalData1_v2.bed.gz

### Code availability

Code used in this study is available at https://github.com/gymreklab/mutea-autosomal.

## Acknowledgements

D.R. was supported by NIH grant GM100233 and is a Howard Hughes Medical Institute investigator. M.G. was supported by NIH/NIMH grant 1U01MH105669-01. Y.E. holds a Career Award at the Scientific Interface from the Burroughs Wellcome Fund. This study was supported by National Institute of Justice grant 2014-DN-BX-K089 (to Y.E. and T.W.) and by a generous gift by Paul and Andria Heafy (Y.E.). We thank Nick Patterson, Mark Daly, Yangyue Wan, and Alon Goren for helpful discussions.

## References

1. Samocha, K. E. et al. A framework for the interpretation of de novo mutation in human disease. Nat. Genet. 46, 944–950 (2014).

2. Petrovski, S., Wang, Q., Heinzen, E. L., Allen, A. S. & Goldstein, D. B. Genic intolerance to functional variation and the interpretation of personal genomes. PLoS Genet. 9, e1003709 (2013).

3. Gulko, B., Hubisz, M. J., Gronau, I. & Siepel, A. A method for calculating probabilities of fitness consequences for point mutations across the human genome. Nat. Genet. 47, 276–283 (2015).

4. M., H. The human functional genome defined by genetic diversity. doi:10.1101/08236.

5. Willems, T. et al. The landscape of human STR variation. Genome Res. 24, 1894–1904 (2014).

6. Mirkin, S. M. Expandable DNA repeats and human disease. Nature 447, 932–940 (2007).

7. Houge, G., Bruland, O., Bjørnevoll, I., Hayden, M. R. & Semaka, A. De novo Huntington disease caused by 26-44 CAG repeat expansion on a low-risk haplotype. Neurology 81, 1099–1100 (2013).

8. Amiel, J., Trochet, D., Clément-Ziza, M., Munnich, A. & Lyonnet, S. Polyalanine expansions in human. Hum. Mol. Genet. 13 Spec No 2, R235–43 (2004).

9. Press, M. O., Carlson, K. D. & Queitsch, C. The overdue promise of short tandem repeat variation for heritability. Trends Genet. 30, 504–512 (2014).

10. Gymrek, M. et al. Abundant contribution of short tandem repeats to gene expression variation in humans. Nat. Genet. 48, 22–29 (2015).

11. Quilez, J. et al. Polymorphic tandem repeats within gene promoters act as modifiers of gene expression and DNA methylation in humans. Nucleic Acids Res. 44, 3750–3762 (2016).

12. Hause, R. J., Pritchard, C. C., Shendure, J. & Salipante, S. J. Classification and characterization of microsatellite instability across 18 cancer types. Nat. Med. (2016). doi:10.1038/nm.419.

13. Ballantyne, K. N. et al. Mutability of Y-chromosomal microsatellites: rates, characteristics, molecular bases, and forensic implications. Am. J. Hum. Genet. 87, 341–353 (2010).

14. Burgarella, C. & Navascués, M. Mutation rate estimates for 110 Y-chromosome STRs combining population and father-son pair data. Eur. J. Hum. Genet. 19, 70–75 (2011).

15. Sun, J. X. et al. A direct characterization of human mutation based on microsatellites. Nat. Genet. 44, 1161–1165 (2012).

16. Weber, J. L. & Wong, C. Mutation of human short tandem repeats. Hum. Mol. Genet. 2, 1123–1128 (1993).

17. Ellegren, H. Heterogeneous mutation processes in human microsatellite DNA sequences. Nat. Genet. 24, 400–402 (2000).

18. Mallick, S. et al. The Simons Genome Diversity Project: 300 genomes from 142 diverse populations. Nature (2016). doi:10.1038/nature1896.

19. Willems, T. et al. Population-Scale Sequencing Data Enable Precise Estimates of Y-STR Mutation Rates. Am. J. Hum. Genet. 98, 919–933 (2016).

20. Li, H., Heng, L. & Richard, D. Inference of human population history from individual whole-genome sequences. Nature 475, 493–496 (2011).

21. Willems, T., Zielinski, D., Gordon, A., Gymrek, M. & Erlich, Y. Genome-wide profiling of heritable and de novo STR variations. bioRxiv 077727 (2016). doi:10.1101/07772.

22. 1000 Genomes Project Consortium et al. An integrated map of genetic variation from 1,092 human genomes. Nature 491, 56–65 (2012).

23. Li, H. Aligning sequence reads, clone sequences and assembly contigs with BWA-MEM. arXiv [q-bio.GN] (2013).

24. Gymrek, M., Golan, D., Rosset, S. & Erlich, Y. lobSTR: A short tandem repeat profiler for personal genomes. Genome Res. 22, 1154–1162 (2012).

25. Sun, J. X. et al. A direct characterization of human mutation based on microsatellites. Nat. Genet. 44, 1161–1165 (2012).

26. Lek, M. et al. Analysis of protein-coding genetic variation in 60,706 humans. Nature 536, 285–291 (2016).

27. Mastushita, M. et al. A Glutamine Repeat Variant of the RUNX2 Gene Causes Cleidocranial Dysplasia. Mol. Syndromol. 6, 50–53 (2015).

28. Shibata, A. et al. Characterisation of novel RUNX2 mutation with alanine tract expansion from Japanese cleidocranial dysplasia patient. Mutagenesis 31, 61–67 (2016).

29. Goodman, F. R. et al. Synpolydactyly phenotypes correlate with size of expansions in HOXD13 polyalanine tract. Proc. Natl. Acad. Sci. U. S. A. 94, 7458–7463 (1997).

30. La Spada, A. R. & Taylor, J. P. Repeat expansion disease: progress and puzzles in disease pathogenesis. Nat. Rev. Genet. 11, 247–258 (2010).

31. Quinlan, A. R. & Hall, I. M. BEDTools: a flexible suite of utilities for comparing genomic features. Bioinformatics 26, 841–842 (2010).

32. Michaelson, J. J. et al. Whole-genome sequencing in autism identifies hot spots for de novo germline mutation. Cell 151, 1431–1442 (2012).

33. Telenti, A. et al. Deep sequencing of 10,000 human genomes. Proc. Natl. Acad. Sci. U. S. A. (2016). doi:10.1073/pnas.161336511.

34. Huang, Q.-Y. et al. Mutation patterns at dinucleotide microsatellite loci in humans. Am. J. Hum. Genet. 70, 625–634 (2002).

35. Haasl, R. J. & Payseur, B. A. Microsatellites as targets of natural selection. Mol. Biol. Evol. 30, 285–298 (2013).

36. Ballantyne, K.N. et al. Toward male individualization with rapidly mutating y-chromosomal short tandem repeats. Hum. Mutat. 35, 1021–1032 (2014).

37. Amos, W., Kosanović, D. & Eriksson, A. Inter-allelic interactions play a major role in microsatellite evolution. Proc. R. Soc. B 282, 20152125 (2015).

38. Garza, J. C., Slatkin, M. & Freimer, N. B. Microsatellite allele frequencies in humans and chimpanzees, with implications for constraints on allele size. Mol. Biol. Evol. 12, 594–603 (1995).

39. Excoffier, L. & Foll, M. fastsimcoal: a continuous-time coalescent simulator of genomic diversity under arbitrarily complex evolutionary scenarios. Bioinformatics 27, 1332–1334 (2011). 453–457 (2015).

40. Helgason, A. et al. The Y-chromosome point mutation rate in humans. Nat. Genet. 47, 453–457 (2015).

41. Poznik, G. D. et al. Punctuated bursts in human male demography inferred from 1,244 worldwide Y-chromosome sequences. Nat. Genet. 48, 593–599 (2016).

